# Sorting of droplets at kHz rates using absorbance activated acoustic sorting

**DOI:** 10.1101/2022.09.13.507732

**Authors:** Esther S. Richter, Andreas Link, John S. McGrath, Raymond W. Sparrow, Maximilian Gantz, Elliot J. Medcalf, Florian Hollfelder, Thomas Franke

## Abstract

Droplet microfluidics allows one to address the ever-increasing demand to screen large libraries of biological samples. Absorbance spectroscopy complements the golden standard of fluorescence detection by label free target identification and providing more quantifiable data. However, this is limited by speed and sensitivity. In this paper we increase the speed of sorting by including acoustofluidics, achieving sorting rates of target droplets of 1 kHz. We improved the devices design for detection of absorbance using fibre-based interrogation of samples with integrated lenses in the microfluidic PDMS device for focusing and collimation of light. This optical improvement reduces the scattering and refraction artefacts, improving the signal quality and sensitivity. The novel design allows us to overcome limitations based on dielectrophoresis sorting, such as droplet size dependency, material and dielectric properties of samples. Our acoustic activated absorbance sorter removes the need for offset dyes or matching oils and sorts about a magnitude faster than current absorbance sorter.

## Introduction

A powerful aspect of droplet microfluidics is the ability to rapidly screen, and sort small volume droplets (pL) from a large population at high throughput. This is required for many applications in biochemistry and molecular biology which also require sensitive screening^1^. Throughput is key for screening and identification of extremely large libraries and for selection of target cells with the desired properties.^2^

Droplets can be actively sorted by microfluidic deflection into a collection channel using electric, acoustic, magnetic, pneumatic or thermal control.^3^ Dielectrophoretic droplet sorting was used by Baret *et al*.^4^ and achieves sorting rates of several kHz ^5^. However, this sorting technique depends on the droplet volume and the dielectric contrast of the drop and the continuous fluid applying voltages up to several kV. Such electric field strengths can potentially harm cells and electroporation or a loss of viability has been reported. ^6,7^

We use acoustic sorting since a low electric field on a piezoelectric substrate is required and therefore no harm to the cells occurs. Acoustic droplet sorting, has shown to sort droplets and particles at kHz droplet rates.^3,7–11^ The sorting is enabled by acoustic streaming inside a channel changing the flow and deflecting droplets. Acoustic sorting is therefore independent of droplet content, properties or volume and uses lower applied power to achieve sorting.^8,9^

Sorting in droplet-fluidics is often triggered by a fluorescence signal of the drop content, for example a fluorescent cell or assay. It is a powerful tool for enzyme activity detection and activated droplet sorting (FADS) uses a laser to excite fluorescence from a droplet. ^3,4,8,9,12^ The presence of a fluorophore label within the droplets is required to produce the fluorescent readout, thus introducing an artefact into the system.^2^ Fluorescent sorting devices usually require expensive and complex optical set-ups^13^.

In contrast, absorbance detection has the potential for smaller and simpler set-ups as well as not requiring a fluorescent label.^13^ Absorption spectroscopy is widely used as a readout in colorimetric assays^14^, protein quantification^15^ and enzyme kinetics^16^ and therefore compliments fluorescent techniques. The standard samples are measured in a cuvette requiring a sample volume of typically 0.2 – 4 mL with 1 cm path lengths^16^. More recently Nano Drop spectroscopy has developed where sample volumes of around 1 µL are used.^17^ However, with all these methods sample measuring rates are low, with each sample taking minutes to measure.^18,19^

Droplet microfluidics offers the possibility of combining the power of absorption spectroscopy with sample volumes in pico-litres and sample throughput in the kHz range.^20^

A challenge with absorbance detection in droplets is their small volume (pL), and the resulting short optical path length (μm), which has an impact on the sensitivity of the detection.^2,14^ Absorbance can be detected using integrated optic fibres. The voltage signals corelate to Beer-Lamberts Law and the droplet solutions absorbance can be calculated.

Optic fibres are easy to integrate into microfluidic chips, they are low cost and flexible leading to simpler over-all set-ups for sorting experiments^13,14^ and have been used for fluorescence as well as absorbance detection in microfluidic chips ^12,13,21^.

Gielen *et al*. ^12^ introduced an absorbance activated droplet sorter (AADS) using dielectrophoresis and optic fibres for sorting of droplet after their absorbance readout with 300 Hz sorting rates. They claim that sorting of concentrations below 100 µM is not possible due to signal artefacts in the droplet caused by the different refractive indices of the oil and water phase. For their directed evolution experiment 1 mM WST1-formazan was added to the assay to mask the artefacts in the signal.^12^ More recently, full UV-Vis spectra screening and sorting has been demonstrated by Probst *et al*.^2^ *and* Duncombe *et al*. ^15^ with achieved droplet rates below 100 Hz since full spectra screening decreases throughput. Duncombe *et al*. ^15^ incorporated an increased path length from 50 to 300 μm and droplet volumes above 300 pL.

Here, we combine the power of rapid screening using surface acoustic wave (SAW) based sorting and with the advantages of absorbance detection. This enables absorbance triggered sorting rates of about an order of magnitude higher than reported by Gielen *et al*.^12^. To increase sensitivity micro-lenses in the form of an optical air cavity in the PDMS have been used to collimate the divergent light emitted from optic fibres and improve the signal. ^13,21^

Combining fibre optic absorbance detection with acoustic droplet sorting enables us to sort with droplet rates up to kHz rates. We can sort droplets without adding an offset dye at concentrations below 100 μM. By mechanically sealing the PDMS channels to the substrate we can reuse our IDT.

## Results and discussion

### Methodology

The design of the acoustic activated absorbance droplet sorter (AcAADS) is shown in Fig. 1. Aqueous droplets are produced in a flow focusing drop-maker device where fluorinated oil flowing from two sides, as the continuous phase, splits the disperse aqueous phase into droplets. The produced droplets are collected and injected into the AcAADS-chip using the droplet inlet. The oil is used to space the droplets and focus them. The AcAADS-chip can be separated into five components. After the droplet re-injection into the chip the droplets are spaced and then the absorbance is measured using two optical fibres. In the detection area the channel is constricted to 50 μm width. The excitation fibre illuminates the sample with 455 nm light, and the detecting fibre detects the change in voltage. An increase in absorbance shows as a decrease in voltage in the signal. The droplets are then hydrodynamically focused before they enter the sorting region and sorting is triggered by an electric pulse from a signal generator applied to a tapered inter digitated transducer (TIDT) that generates a traveling surface acoustic wave (TSAW). An IDT generates a TSAW which propagates along the surface of the piezoelectric substrate causing internal streaming of the fluid within a channel. The IDT is positioned behind the focusing channels and generates TSAW deflecting the droplet.

**Fig. 1.**
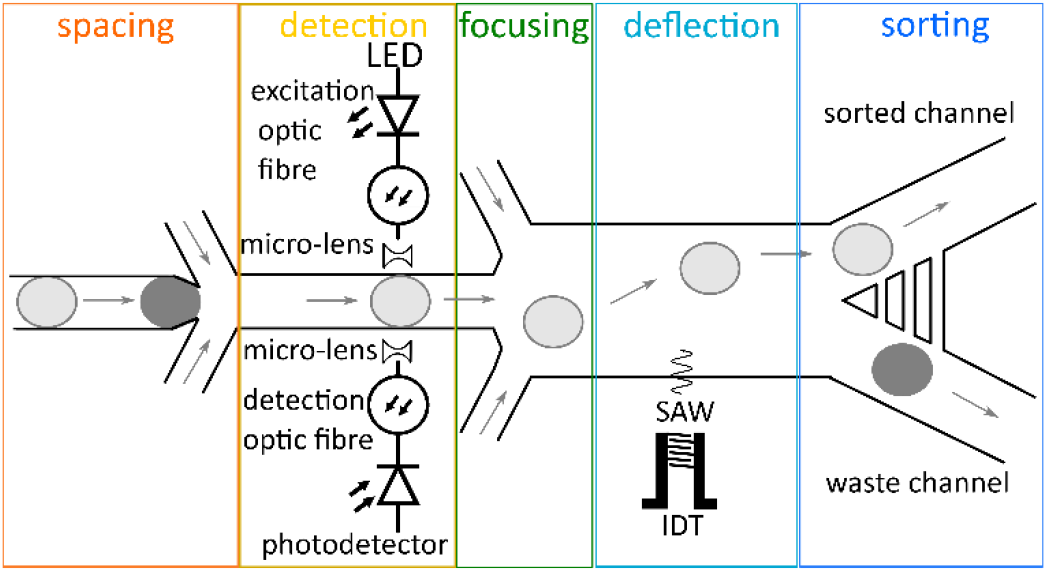
The AcAADS can be separated in the five functional regions spacing of the droplet, absorbance detection, focusing of the droplet to the preferential flow, deflection of the droplet by SAW and sorting into the two waste outlet channel. The arrows indicate flow directions and the the colour of the droplets reperesent different absorbance values of the droplet content.

### Absorbance signal detection

We probe the absorbance detection using the food dye Tartrazine as the absorbing medium and record the signals as peaks of droplets passing through the detection region of the chip. Tartrazine droplets in fluorinated oil have a characteristic peak signal profile. For 30 μM Tartrazine concentration the signal is composed of two downwards peaks, below the oil baseline, and an upwards peak in between, above the baseline as shown Fig. 2A. This is a typical profile observed for concentrations below 100 μM. For higher concentrations the whole peak profile is below the oil baseline (data not shown here). This characteristic signal shape has also been reported by other groups, Gielen *et al*.^12^ and Hengoju *et al*.^13^, and they suggest that the mismatch of refractive index of the oil, *n*^_*oil*_^ = 1.287, and water-phase, *n*^_*water*_^ = 1.33, causes these artefacts. The peaks below the baseline are generated by scattering and refraction at the front and rear edges of the passing droplets.^2^ Gielen *et al*.^12^ claim that these effects mask the absorbance signal of the droplets.

**Fig. 2.**
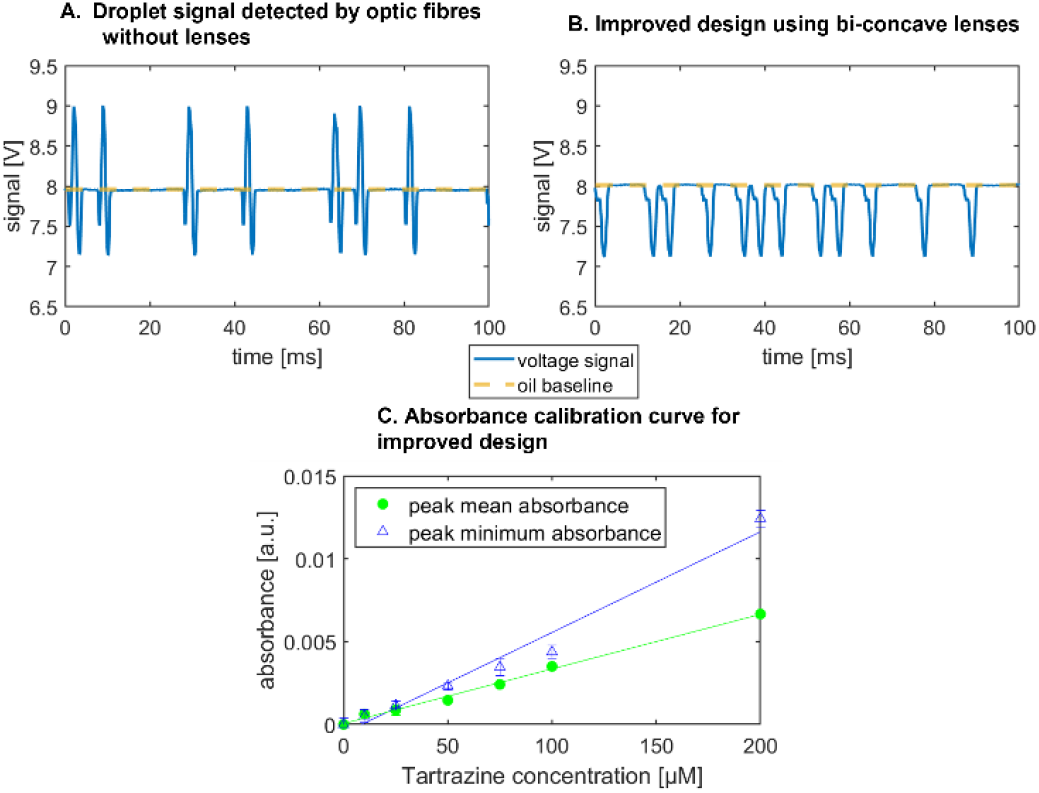
**A** Signal of seven 30 μM Tartrazine droplets passing the detector of the AcAADS-chip. The signal baseline of the oil was set to 8 V of the LED. The two downward peaks of a single droplet signal profile represent the entry and exit of a droplet in the region of detection, respectively. **B** Signal of twelve 30 μM Tartrazine droplets using the improved design with integrated bi-concave micro-lenses. The signal for a single drop is entirely below the baseline and the central peak has been diminished to become a small shoulder in the profile. **C** The average of the mean and minimum absorbance values of 1500 droplets is plotted against the Tartrazine concentration. Absorbance *A* is calculated using Beer-Lamberts law from the measured droplet voltage signals, using the water droplet signal as reference. The droplet mean value for a single droplet is determined by the sum of the voltage signal, taken every 5 ns, divided by the peak duration. The linear regression function for the absorbance data of the tartrazine concentration is *f*(*x*) = *m x* + *b* with *m*_*mean*_ = 3.29 · 10^−5^ ± 1.048 · 10−6, *b*_*mean*_ = 5.21 · 10^−5^, *m*_*min*_ = 6.08 · 10^−5^ ± 4.48 · 10^−6^ and *b*_*min*_ = − 5.32 · 10^−4^ respectively.

To increase the sensitivity and reduce the artefacts of the signal we present an improved chip design. We integrate micro-lenses into the PDMS mould to collimate the illuminating light and increase the yield of transmitted light collected by the detection fibre. We use lenses made of cavities in the PDMS on each side of the microfluidic channel. One lens is used to collimate the light before passing the channel and evenly illuminate the droplet. The second lens on the backside of the channel collects the light and focuses it into the detection fibre, as shown in Fig. 3. Since the medium of the lens is air with a refractive index *n*_*air*_ = 1 and the refractive index of the environment is cured PDMS is larger, *n*_*PDMS*_ = 1.4, converging lenses are concave.^13^ For compact lens design we use bi-concave lenses.

**Fig. 3.**
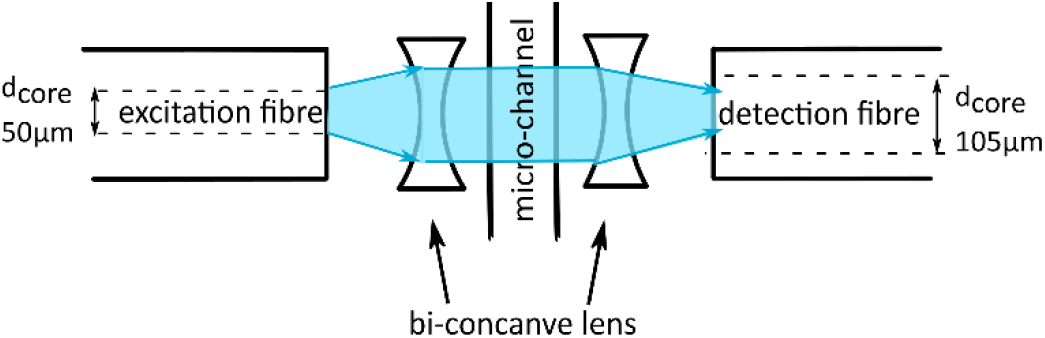
Drawing of the detection area of the chip showing the design of the bi-concave micro-lenses and the collimated light beam (blue). The two lenses have different focal length due to the core diameter of the detection fibre being larger.

Micro-lenses are designed using the thin lens maker’s equation, relating the focal length *f*_*lens*_ with the refractive indices *n* and radii of curvature *r* of the lens surfaces, as given in eq. 1^13,21^. For bi-concave, thin lenses, this equation reduces to

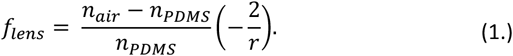

In the new design of the micro-lenses the mechanical stability of the PDMS walls separating the lenses from the micro-channel and the optic fibres need to be considered. If the walls are too thin capillary effects in the fibre insertion guides as well as pressure effect in the microfluidic channel can deform the lenses. With a wall thickness of 100 μm and numerical aperture, *NA* = 0.22, of the fibres we calculate the focal length and determine the curvature of the lenses from eq. 1, see also material section.

The optical improvements using lenses show a cleaned-up signal with reduced scattering and refractive artefacts. The droplet signal profile using the improved optical detection is below the oil baseline and the peak in the middle is diminished and appears only as a shoulder in the decreasing flank of the signal, as shown in Fig. 2 B.

We quantify the signal by determining the absorbance from the voltage signal of the photodetector using the Beer-Lamberts law.

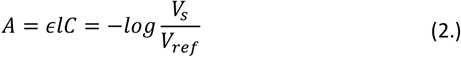

The absorbance *A* is equal to the logarithmic relationship of the sample voltage *V*_*s*_ and the reference voltage *V*_*ref*_, as well as the product of the path length *l*, Tartrazine concentrations *C*_*T*_ and molar extinction coefficient *ϵ*.22 The absorbance is linearly proportional to the concentration of the sample. ^23,24^

We calibrate the system using different aqueous dilutions of Tartrazine (0 - 200 μM) in droplets of the same size and use the signal of water droplets as *V*_*ref*_.

The sorting software generates the mean and minimum peak values for each droplet signal. We calculated the absorbance from these voltages and depicted their linear relation with increasing Tartrazine concentration in Fig. 2 C. Both the mean and minimum values can be used to trigger the sorting decision. The gradient of the peak minimum absorbance values is larger than the mean peak gradient, however the deviation for each concentration is larger. The calibration sensitivity *s*_c_ is given by the slope *m* 25 in the graph in figure 2 C to be *s*^_*c*_^ = *m* = 6.08 ∙ 105 μM^-1^ for the peak minimum data. The minimum distinguishable analytical absorbance signal *A*_*min*_ can be estimated to be *A*_*min*_ = *k* ∙ *σ*_*water*_ with the standard deviation of the water droplet measurements *σ*_*water*_ and *k* being a constant describing the confidence interval. For a typical value of *k* = 3, we obtain *A*_*min*_ = 0.00263. Using the values from figure 2 C the concentration limit *c*_*l*_ = *A*_*min*_/*m* can be calculated to be *A*_*min*_ = 43.2 μM.

In figure 2 C the deviations of droplet signals for detected concentrations with 50 μM difference do not overlap.Whereas, with lower concentration differences there is potential for overlap of the two populations signals.

### Droplet deflection by SAW

When the voltage signal of the droplet crosses the set threshold the SAW is triggered after a set delay time. The delay time is the time the droplet needs to move from the detection area to the point deflection by of SAW. The change in streaming caused by the SAW pushes the droplet into the upper channel as depicted in the droplet trajectory in Fig. 4.

**Fig. 4.**
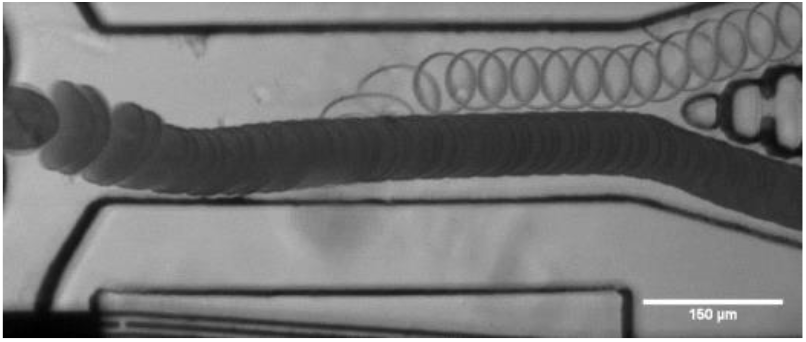
Overlay of frames indicating the trajectory of water and bromophenol blue droplets, where the sorting value is set for the water droplet signal voltage. The dark bromophenol blue droplets are not sorted and flow into the waste channel. The flow is adjusted using the two focusing inlets. When a water droplet is detected, the SAW is triggered after a set delay time and the droplet is deflected into the upper channel.

For tapered IDTs the point of actuation of the SAW is frequency dependent. It is important to set the correct delay time for optimal droplet deflection for successful sorting. If the delay time is not correct the deflection might not be enough for sorting, or the droplet might even be pushed down which is caused by the two vortexes of the SAW. We deflected droplets with the SAW on constantly, using different SAW powers and plotted their trajectories shown in Fig. 5. With the different applied powers, the droplets are deflected at the same location, but to different heights. The higher the power the greater the deflection. The droplets are firstly being pulled down by the SAW vortices. The point of deflection for sorting should be chosen where the slope is positive again. The measured distance of this point in the trajectories is 245 μm in the x direction to the right of the beginning of the IDT aperture and is fixed for this device channel geometry and IDT. We can deflect droplets within 100 μm of the SAW actuation location where the trajectory turns negative again. When the droplet velocity is known the pulse duration can be derived from the distance of the actuation and turning off point. The pulse duration further affects the droplet spacing. A too long pulse duration can change the trajectory of the following drop leading to potential false positive sorting events.

**Fig. 5.**
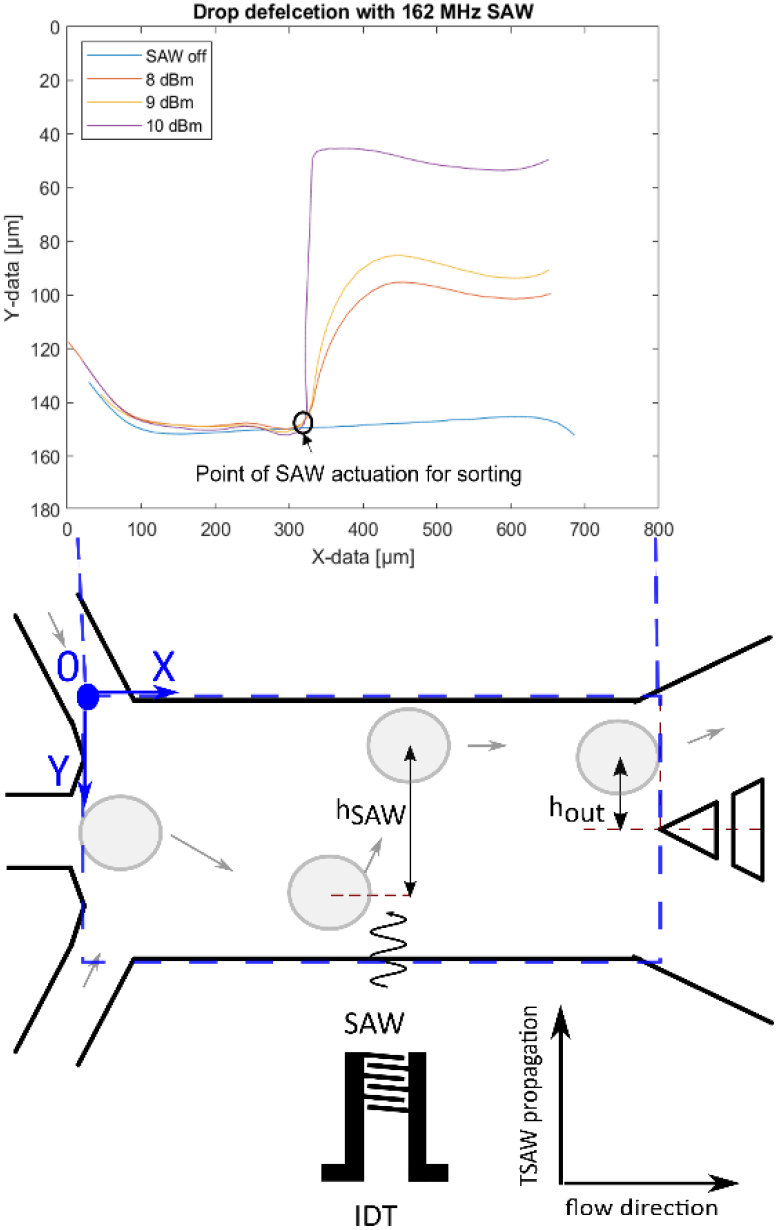
Graph showing the trajectories of droplets for different applied powers when the SAW is on constantly. This is used to establish the active area of the SAW and the optimal droplet position in relation to the IDT for deflection. But also, the optimal switch on time of the SAW. For optimal deflection the droplet should be at the point of the trajectories where the slope becomes positive. Droplets are deflected within a short distance, ca. 100 μm.

To characterize the deflection of droplets in the AcAADS device droplets of the same size (70 μm diameter) where deflected with a SAW frequency of 162 MHz at three different droplet rates (300 Hz, 800 Hz and 1300 Hz) at 251 mW power and for pulse durations ranging from 50 μs and 1 ms. The deflection height was measured using ImageJ after the SAW actuation (point) *h*_*SAW*_ and at the tip of the separating pillars at the two outlet channels *h*_*out*_, as illustrated in Fig. 5.

The Graphs in Fig. 6 A and B show the droplet deflection *h*_*SAW*_ and *h*_*out*_ for the three droplet rates and pulse length range. With an increase in pulse duration the deflection height increases. Also, the droplet rate influences the deflection, for lower droplet rates the defection is larger. If the deflection at the outlet channel *h*_*out*_ is negative, then the deflection is not sufficient to push the droplets into the upper sorting channel and thus no sorting occurs. Also, if *h*_*out*_ is not larger than the radius of the droplet it hits the separation post and may split. The sorting efficiency for the deflection experiments is plotted in the graph in Fig. 6 C. For higher droplet rates a longer SAW pulse is needed for higher sorting efficiency. If *h*_*out*_ is at or close to zero, the droplets flow into either outlet channel. Some of the deflected droplets are sorted and some are not, as shown in the experiment of 1300 Hz and a pulse length of 200 μs. In case of the 1300 Hz and 1 ms experiment, the pulse length was too long, causing the SAW triggered by a following droplet to affect the prior droplet. This affects the sorting efficiency as some droplets are not sorted as they are being pushed down by a following triggered SAW. Droplet rates higher than 1.5 kHz are possible, but the droplets, at the size of 70 μm diameter, form a bullet shape in the squeezed detection area. When they enter the wider sorting region the corners of droplets, in the bullet shape, can split as small satellite droplets. Therefore, we show sorting experiments above 1 kHz, but below 1.5 kHz, where this does not happen.

**Fig. 6.**
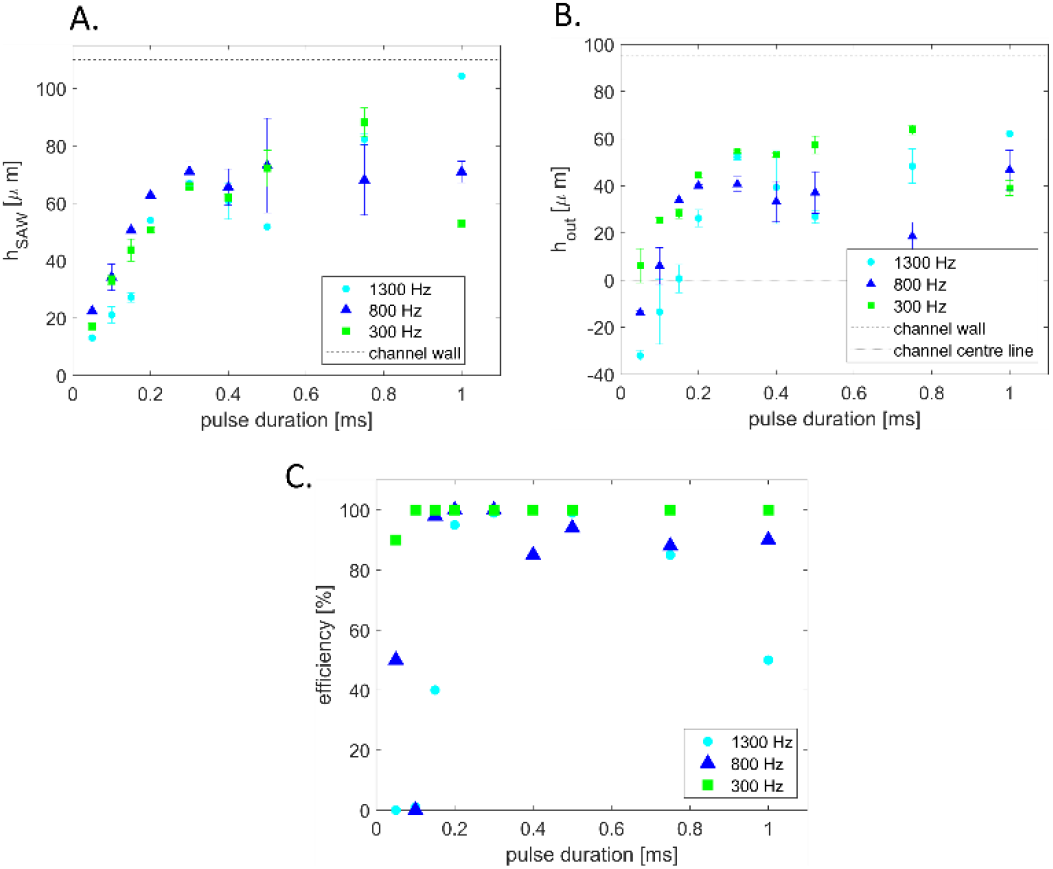
Graphs showing the deflection of 70 μm droplets using 251 mW for pulse lengths ranging between 50 μs and 1 ms and three droplet rates (300 Hz, 800 Hz and 1300 Hz). **A** droplet deflection at the SAW active region h_SAW_ and **B** at the beginning of the outlet channels h_out_. **C** sorting efficiency for 1300 Hz, 800 Hz and 300 Hz droplet rates at 251 mW applied power for different pulse length ranging between 50 μs and 1 ms.

### Sorting droplets with 30 μM and 80 μM Tartrazine dye

To validate the sorting process, we demonstrate the correlation between droplet signal detection and droplet sorting. Therefore, our sorting software enables simultaneous recording of both the detected absorbance signal and the video of the droplet flowing in the microchannel. We then match both records after the experiment frame by frame by synchronizing their time axis using the frame rate and a mutual time stamp.

Prior to sorting, we need to adjust the sorting parameters for signal threshold and flow conditions. Therefore, we observed the signal of the two populations of Tartrazine droplets, 30μM and 80μM and display them in a scatter plot. In the scatter plot of the peak values two populations are distinguishable. We set the sorting threshold between the two populations at 7,41V. Droplets with minimum peak values below 7.41 trigger the acoustics and are deflected into the sort outlet channel. Then, we determine the time it takes from the interrogation to the sorting area by monitoring the droplet flow and set the time delay td= 1.12ms. This time corresponds to the time span between detecting a absorbance signal above the threshold and triggering the droplet sort signal to activate the IDT.

Using the overlay of the droplet signal and sorting video we can identify each target droplet to be above the threshold and confirm corresponding droplet sorting in the designated outlet with the video as shown in Fig. 7.Here we demonstrate a sorting experiment of two droplet populations with Tartrazine concentrations of 30μM and 80μM at 1kHz rate.

**Fig. 7.**
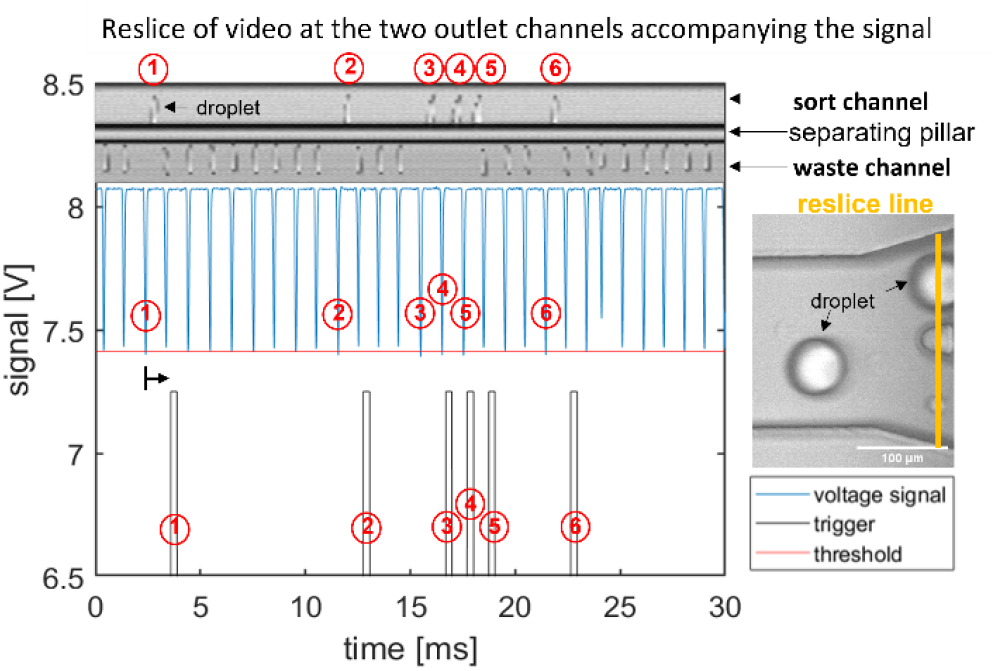
Graphs demonstrating detection and sorting process of 30μM and 80μM Tartrazine droplets at 1 kHz sorting rate. The upper micrograph is a spatiotemporal representation of the sorting process (reslice image). The image is calculated from pixels taken at a fixed line frame by frame from the droplet sorting video and then plotted against time. We position the reslice line across the the two outlets as highlighted itin orange in the micrograph (right image). Hence, the upper channel in the reslice micrograph is the sort channel with 6 deflected droplets, and the lower is the waste channel containing 24 not deflected drops. The original droplet absorbance signal data is shown in the central plot (blue curve). It is analysed in real time using our LabVIEW software generating the droplets minimum and mean peak values for each drop. When the minimum peak value crosses the threshold (red) a sorting trigger signal is generated (bottom plot in grey). The trigger signal is triggered (grey) with the pre-set time delay of *t*_delay_=1.12ms, as indicated by the black arrow that accounts for the time it takes the drop to flow from detection to sorting area. Individual target droplets (80μM Tartrazine) are labelled by numbers in the detection, trigger and resliced images for comparison. The figure covers an extract of 30ms take from a 1s sorting process.

Smaller concentration differences the absorbance signals of both populations are not clearly separated and overlap in the scatter plot, and sensitivity and specificity start to decrease.

## Conclusion

In this paper we present an acoustic activated absorbance droplet sorter with integrated bi-concave lenses to focus the light beam. We reduce the scattering effect reported earlier and increase sensitivity. The resulting droplet signal shape enables us to sort between droplets with concentration difference of 50 μM, with smaller concentration differences the cluster in the peak scatter plots are overlapping. Furthermore, we can sort droplets with 70 μm diameter at 1 kHz droplet rates efficiently faster than the previously reported 300 Hz. We show sorting experiments of droplet of two droplet populations with 30 and 80 μM. To further increase the droplet rate the channel design needs to be adjusted to match the drop size and to reduce the droplet deformation that eventually causes drop disintegration and the formation of daughter droplets. For higher sorting rates higher SAW powers might need to be applied for successful sorting as a longer acoustic pulse length may affect neighbouring droplets. The separation of the optofluidic components, all integrated into the PDMS mould, and the piezoelectric chip that is in purely mechanical contact, allows us to reuse the chip and operate multiple different channels on one chip. Medcalf et al.^26^ use refractive index matching and improved micro-channels to increase the absorbance signal quality and throughput for dielectrophoretic droplet sorting. Combining both techniques has the potential for further improvements for absorbance droplet sorting.

## Material and methods

### Microfluidic-chip fabrication

The AcAADS-chip consists of a tapered IDT (TIDT), PDMS micro-channels and optic fibres. The TIDT generates traveling surface acoustic waves with a resonance frequency f of 161-171 MHz. It consists of two comb-like gold electrodes on LiNbO3 (128°rot-Y-cut). The electrodes have 60 finger pairs, are 500 μm long and due to the tapered design their periodicity varies from 23.0 to 24.4 μm.^8,9^

Metal deposition is used to deposit gold onto the LiNbO3 substrate and S1818 photolithography generates the IDT’s structure as described in Link *et. al*^27^.

Microfluidic channels were fabricated using standard two-layer SU8-photolithography and PDMS. A two-layer photolithography was needed to generate the height for the optic fibre insertion. The fluidic channels are 80 μm and the insertion guides are 130 μm in height respectively.

The channel structures in PDMS are generated using replica moulding. PDMS base (Sylgard 184 Silicone Elastomer, Sigma Aldrich, USA) and curing agent mixed in a 10:1ratio and poured on the wafer in a Petri dish and cured at 75°C for 4 hours.

The cured PDMS was cut into the single chips, triangle openings are cut at the fibre insertion channels and a 1.0 mm biopsy puncher (Miltex Biopsy Punch, Kai Medical, Japan) used to create the inlets and outlets.

The PDMS was aligned on the IDT and pressed onto the LiNbO_3_ substrate, and the channels sealed using a custom-made chip holder. This makes the IDT reusable, as the PDMS can be removed and, after cleaning the surface, a new chip can be aligned onto it.

The electrode pads of the IDT are aligned with the PCB connections and conductive (silver) paint (RS Components, UK) creates a connection between the chip and the PCB.

### Optofluidic components

Multimode optical fibres of two different core diameters are used, the illumination fibre’s core diameter is 50 μm (M14L, Thorlabs, USA) and the detection fibre has a core diameter of 105 μm (M15L, Thorlabs, USA). The illumination fibre is inserted into the cavity guide structure next to the IDT. PDMS was injected to fill the remaining air cavity of the guide structure. This stabilizes the fibres after the PDMS is cured avoiding a large gradient in refractive index constant. The detecting signal is analysed by a LabVIEW software that is also used for the sorting. We designed the lens to collimate the emitted light cone that enters the lens. The cone of light is dependent on the numerical aperture *NA* and the fibre core diameter. To calculate the radii of curvature we set the focal length *f*_*lens*_ as the PDMS wall thickness, 100 μm, and the distance from the approximated focal point of the cone of light using the aperture angle *Θ* to the fibre tip. Depending on the individual chip the position of the collecting fibre for the improved signal can vary. The angle of aperture is also used to calculate the beam diameter at the lens. To ensure all the light is collected by the lens we set the lens diameter larger than the calculated beam diameter to minimum 150 μm. We calculated a radius of curvature for the illuminating lens of 147 μm and 245 μm for the focusing lens.

### Experimental set-up

For recoding the videos, we use an inverted microscope (IX73, Olympus, Japan) and a fast camera (Fastcam Mini UX50, Photron, Japan). A custom-made pressure system is used to generate all flows inside the chip. The optical fibres are connected accordingly, the 105μm fibre (M15L, Thorlabs, USA) to the photodetector (PDA100A2, Thorlabs, USA) and the 50μm (M14L, Thorlabs, USA) with the LED, 455nm wavelength emission, (M455F1, Thorlabs, USA). The photodetector detects a voltage signal and an increase in absorbance causes a decrease in voltage. The photodetector connects to an FPGA card NI PCIe-7841R (National instruments, USA) and the whole system is controlled by the LabView software.

A switch (ZX80-DR230-S+, Mini-circuit, USA) connects the FPGA card with a signal generator (SMB100A, Rohde&Schwarz, Deutschland) and the PCB of the AcAADS-chip. If a droplet signal crosses a pre-set threshold the SAW is triggered and sorts the target droplet. In the LabVIEW software the parameters for the sorting threshold, the delay time and pulse length are set accordingly.

### Droplet production

The droplets of 70 μm diameter are produced in a droplet-maker-chip using flow focusing (420 μL/hr oil, 65 μL/hr water-phase) using syringe pumps (PHD ULTRA, Harvard Apparatus, US) and collected in a prepared collector pipette tip.

HFE Novec 7500 (3M, USA) fluorinated oil with 1.8% FluoSurf surfactant (Emulseo, France) was used for the continuous phase with the aqueous solutions as the dispersed phase forming the droplets. Surfactants are used to stabilize the droplet interface and prevent bursting or merging of droplets while re-injecting them into a chip. To analyse the detection capabilities Tartrazine dilutions (Sigma Aldrich, USA) were used and for the deflection characterization experiments, a population mix of 0.03 and 1 mM Tartrazine was used. In the microscope videos the bromophenol blue droplets are visibly darker than the water ones, thus enabling visible confirmation of the sorting efficiency.

### Experimental methods

The flow in the channel and droplet reinjection were controlled using the pressure system. The baseline was set for HFE oil by applying 8 V to the LED. For each droplet content concentration three signals over 1 s, and histograms containing the mean and minimum voltage of 1500 droplets are saved using the sorting software.

The deflection of droplets inside the AcAADS-chip was characterized using 0.03 and 1 mM Tartrazine droplets reinjected into the chip at three different droplet rates (300, 800 and 1300 Hz), see Table 1 for the applied pressures. For the acoustics we applied 251 mW (24 dBm) and 162 MHz and varied the pulse duration from 50 μs to 1 ms (see Fig. 6).

**Table 1.** The table provides the applied pressures to generate the flows for the three droplet rates of the droplet deflection length experiments.

| drop rate<br>[Hz] | sample<br>reinjection<br>[mbar] | spacing<br>[mbar] | sheath<br>flow 1<br>[mbar] | sheath<br>flow 2<br>[mbar] |
| --- | --- | --- | --- | --- |
| 300 | 40 | 33 | 40 | 10 |
| 800 | 60 | 50 | 50 | 10 |
| 1300 | 90 | 80 | 70 | 10 |

For each parameter set, three videos were taken, and 10 droplets for the deflection were measured to calculate the average deflection. The deflection was measured, from the droplet centre point, using ImageJ. The sorting efficiency represents the percentage of successfully sorted droplets from the droplet population with a signal crossing the set threshold.

For the sorting experiments with droplet concentrations below 100 μM two droplet populations of 30 and 80 μM Tartrazine, were produced and mixed in the reinjection tip. The flows inside the AcAADS were adjusted by setting the pressures (150 mbar applied on droplet reinjection, 140 mbar spacing, 80 and 10 mbar focusing). Once the droplet rate is set, the delay time (*t*^_*delay*_^ = 1.12 ms) is calculated and the sorting threshold set 7.41 V. Droplets with a minimum peak below the threshold are sorted. The SAW parameters set to 162 MHz frequency, 251 mW (24 dBm) power and 300 μs pulse duration. Simultaneous saving of the video and signal is triggered, and sorting efficiency is verified by overlaying the signal and the video.

## Acknowledgements

This work is part of a project that has received funding from the European Union’s Horizon 2020 research and innovation programme under the Marie Skłodowska-Curie grant agreement No. 813786 (EVOdrops). This work reflects only the author’s view, and the Research Executive Agency is not responsible for any use that may be made of the information it contains.

## References

1 S. Neun, P. J. Zurek, T. S. Kaminski and F. Hollfelder, Ultrahigh throughput screening for enzyme function in droplets, Elsevier Inc., 1st edn., 2020, vol. 643.

2 J. Probst, P. Howes, P. Arosio, S. Stavrakis and A. Demello, Anal. Chem., 2021, 93, 7673–7681.

3 H. D. Xi, H. Zheng, W. Guo, A. M. Gañán-Calvo, Y. Ai, C. W. Tsao, J. Zhou, W. Li, Y. Huang, N. T. Nguyen and S. H. Tan, Lab Chip, 2017, 17, 751–771.

4 J. C. Baret, O. J. Miller, V. Taly, M. Ryckelynck, A. El-Harrak, L. Frenz, C. Rick, M. L. Samuels, J. B. Hutchison, J. J. Agresti, D. R. Link, D. A. Weitz and A. D. Griffiths, Lab Chip, 2009, 9, 1850– 1858.

5 A. Sciambi and A. R. Abate, Lab Chip, 2015, 15, 47–51.

6 P. A. Garcia, Z. Ge, J. L. Moran and C. R. Buie, Sci. Rep., 2016, 6, 1–11.

7 R. Zhong, S. Yang, G. S. Ugolini, T. Naquin, J. Zhang, K. Yang, J. Xia, T. Konry and T. J. Huang, Small, 2021, 2103848, 1–9.

8 L. Schmid, D. A. Weitz and T. Franke, Lab Chip, 2014, 14, 3710–3718.

9 T. Franke, S. Braunmüller, L. Schmid, A. Wixforth and D. A. Weitz, Lab Chip, 2010, 10, 789– 794.

10 D. J. Collins, A. Neild and Y. Ai, Lab Chip, 2016, 16, 471–479.

11 S. Li, X. Ding, F. Guo, Y. Chen, M. I. Lapsley, S. C. S. Lin, L. Wang, J. P. McCoy, C. E. Cameron and T. J. Huang, Anal. Chem., 2013, 85, 5468–5474.

12 F. Gielen, R. Hours, S. Emond, M. Fischlechner, U. Schell and F. Hollfelder, Proc. Natl. Acad. Sci. U. S. A., 2016, 113, E7383–E7389.

13 S. Hengoju, S. Wohlfeil, A. S. Munser, S. Boehme, E. Beckert, O. Shvydkiv, M. Tovar, M. Roth and M. A. Rosenbaum, Biomicrofluidics,, DOI:10.1063/1.5139603.

14 R. M. Maceiczyk, D. Hess, F. W. Y. Chiu, S. Stavrakis and A. J. DeMello, Lab Chip, 2017, 17, 3654–3663.

15 T. A. Duncombe, A. Ponti, F. P. Seebeck and P. S. Dittrich, Anal. Chem., 2021, 93, 13008– 13013.

16 S. M. Nilapwar, M. Nardelli, H. V. Westerhoff and M. Verma, Absorption spectroscopy, Elsevier Inc., 1st edn., 2011, vol. 500.

17 M. L.C. Passos and M. L. M.F.S. Saraiva, Meas. J. Int. Meas. Confed., 2019, 135, 896–904.

18 A. Micsonai, F. Wien, L. Kernya, Y. H. Lee, Y. Goto, M. Réfrégiers and J. Kardos, Proc. Natl. Acad. Sci. U. S. A., 2015, 112, E3095–E3103.

19 H. Westerhoff, M. Verma and D. Jameson, Methods in Systems Biology, Elsevier Science, 2011.

20 T. Yang, S. Stavrakis and A. DeMello, Anal. Chem., 2017, 89, 12880–12887.

21 I. Rodríguez-Ruiz, T. N. Ackermann, X. Muñoz-Berbel and A. Llobera, Anal. Chem., 2016, 88, 6630–6637.

22 T. G. Mayerhöfer, H. Mutschke and J. Popp, ChemPhysChem, 2016, 1948–1955.

23 M. F. Vitha, Spectroscopy: Principles and Instrumentation, Wiley, 2018.

24 K. Wilson and J. Walker, Principles and Techniques of Biochemistry and Molecular Biology, Cambridge University Press, 2010.

25 A. J. Demello, ACS Sensors, 2022, 7, 1235–1236.

26 E. J Medcalf et al., Ultrahigh-throughput Absorbance Activated Droplet Sorting (UHT-AADS) for enzyme screening at kilohertz frequencies. bioRxiv (2022).

27 A. Link, J. S. McGrath, M. Zaimagaoglu and T. Franke, Lab Chip, 2022, 22, 193–200.

